# Efficacy of vaccines against *Flavobacterium* infections in fish: A systematic review and meta-analysis

**DOI:** 10.64898/2026.09.11.750515

**Authors:** Yuanwei Geng, Zulqarnain Baqar, Ruoxi Zhu, Yongtao Zhu, Haitham Mohammed, Ibrahim Elsohaby, Wenlong Cai

## Abstract

*Flavobacterium* species cause substantial economic losses in aquaculture, but no quantitative synthesis of vaccine efficacy across species, hosts, and platforms has been evaluated yet. This systematic review and meta–analysis aimed to evaluate the efficacy of vaccines against *Flavobacterium* infections in fish. We systematically searched PubMed, Web of Science, and Scopus for controlled challenge trials. The pooled risk ratio was calculated using a random effects model. Heterogeneity was assessed, and publication bias was evaluated by funnel plot and Egger’s test. Sixty studies were included in this study, 59 of which were included in the main meta-analysis (377 comparisons) after screening. The overall pooled risk ratio was 0.572 (95% confidence interval: 0.55–0.60), corresponding to a relative percent survival of 42.8%, with high heterogeneity. Efficacy differed significantly by pathogen species, fish host, vaccine type and administration route. The highest protection by pathogen species was against *F. davisii* in channel catfish (relative percent survival = 86.8%), and the lowest against *F. columnare* in grass carp (relative percent survival = 30.6%). Live attenuated vaccines induced the highest protection, followed by inactivated and subunit vaccines. Among administration routes, immersion and oral vaccination achieved comparable protection, both outperforming injection. Significant publication bias was detected (*P*<0.001). Vaccination effectively reduces fish mortality from *Flavobacterium* infections, but its efficacy is highly context-dependent. Taken together, standardized challenge protocols and improved reporting are needed. Priority should be given to developing effective vaccines against *F. oreochromis* and *F. columnare*.

## 1 Introduction

The genus *Flavobacterium* includes several major bacterial pathogens in aquaculture, among which *F. psychrophilum* and the columnaris-causing bacteria (CCB) are recognized as the two most detrimental ^1–3^. *F. psychrophilum* primarily infects salmonids such as rainbow trout (*Oncorhynchus mykiss*) and Atlantic salmon (*Salmo salar*), causing bacterial coldwater disease (BCWD) and rainbow trout fry syndrome (RTFS) ^4–6^. Outbreaks typically occur at water temperatures below 15 °C, and both clinical signs and mortality are strongly dependent on fish size ^7,8^. In juvenile salmonids, *F. psychrophilum* causes an acute, systemic infection (RTFS), which is characterised by rapid bacterial dissemination and multi-organ colonization, resulting in high mortality ^4,9,10^. In older or larger fish, the disease typically presents as ulcerative and necrotic skin lesions, most prominently on the caudal peduncle (peduncle disease), posing a significant threat to salmonid aquaculture ^5,8,11,12^. The term “columnaris-causing bacteria” (CCB) represents the four distinct species, *F. columnare*, *F. covae*, *F. davisii*, and *F. oreochromis*, that were reclassified from the different genomovars of *F. columnare* by LaFrentz et al. based on genetic differences ^13,14^. Although these four species differ in virulence and host specificity, they are all causative agents of columnaris disease ^15,16^. Columnaris disease affects a broad range of freshwater fish species and typically occurs at water temperatures above 20°C ^3^. However, columnaris disease has also been reported in coldwater species such as trout at water temperatures as low as 12–14 °C ^17^. In addition to the major pathogens mentioned above, *Flavobacterium johnsoniae* and *Flavobacterium araucananum* are also recognized as fish-associated flavobacteria ^18,19^, though they are not generally regarded as major pathogens. *F. araucananum* was first isolated from diseased salmonids in South America and later recovered from fish in North America ^19,20^; its pathogenic mechanisms remain unclear. *F. johnsoniae* is an opportunistic species that has been isolated from superficial lesions of farmed salmonids and other fish ^20,21^. Experimental bath infections induced fin rot and skin ulcers with systemic spread ^18^; high-dose injection caused mortality in zebrafish and common carp ^22,23^. These diseases impose substantial economic losses on global aquaculture. The efficacy of antibiotics against *Flavobacterium* species is increasingly challenged by antimicrobial resistance (AMR), with many isolates exhibiting resistance to multiple drugs ^24–26^. This trend stresses the urgent need for alternative prophylactic measures. Vaccination has become the most practical and sustainable approach for disease prevention in aquaculture ^27,28^. Over the past two decades, the development of *Flavobacterium* vaccines has expanded from conventional inactivated bacterins to a variety of products. Inactivated vaccines, mostly prepared by formalin treatment in the early stage, were administered by injection, immersion, or oral routes. Injectable inactivated vaccines often rely on oil-based adjuvants, such as mineral oil or Montanide formulations, to enhance protective efficacy ^29,30^; in the absence of adjuvants, protection is often ineffective or variable ^31,32^. Currently, protective immunity can be achieved without oil adjuvants when inactivated vaccines are administered via immersion formulated with nanoparticle carriers (e.g., chitosan or cationic lipids) ^33–35^. Oral vaccination with inactivated bacterins via feed induced only partial protection ^36^. Live attenuated vaccine candidates, generated by passage on increasing concentrations of rifampicin or by targeted gene knockout, have been developed primarily against *F. psychrophilum* and *F. covae* ^37–40^. The protective efficacy of such attenuated strains can be further enhanced by cultivation under iron-limited conditions ^41^. Subunit vaccines based on immunogenic proteins represent another major approach, with protective antigens including HCD, AtpD, and GdhA in *F. psychrophilum* ^42^; GldJ in *F. columnare* ^43^; and DnaK in *F. covae* ^44,45^. These subunit vaccines are typically administered by injection or immersion; however, most require oil-based adjuvants to induce protective immunity ^45,46^. When delivered orally, vaccines formulated with alginate microencapsulation or feed-based delivery have provided only partial and less consistent protection ^36,47^. Polyvalent and cross-protective vaccines have also been explored, including polyvalent inactivated vaccines containing multiple serotypes ^30,48^, bivalent nanovaccines targeting *F. oreochromis* with *Francisella orientalis* or *F. covae* with *S. iniae* ^49,50^, and live attenuated vaccines that cross-protect against *F. columnare* and *F. covae* ^39,51^.

Despite the accumulation of vaccine efficacy data over the past two decades, direct comparisons across trials remain difficult because of substantial heterogeneity in challenge protocols, vaccine formulations, and outcome reporting. Studies vary in which antigens are selected, whether boosters are given, how outcomes are measured, and how long mortality is monitored, with observation periods spanning from 10 to 28 days ^29,52^. Some trials report cumulative mortality or survival rate ^46,53^, others report relative percent survival (RPS) ^54,55^, and several provide only immunological readouts that cannot be translated into quantitative estimates of protection ^56^. This also means that when two trials report different levels of protection, it is often impossible to determine whether the difference reflects a true effect of the vaccine or simply a variation in study design. For example, two recombinant heat shock protein vaccines were tested against *F. columnare* in channel catfish. One was based on DnaJ and delivered by injection with Freund’s complete adjuvant; the other was based on DnaK and delivered by immersion without adjuvant. Both elicited specific antibody responses, but only DnaK conferred significant protection ^44,45^. The DnaJ vaccine did not protect despite the use of an adjuvant. Antigen choice, delivery route, and adjuvant use all differed, and the factor responsible for the difference in protection cannot be identified from these two trials alone. Against this background, a systematic review and meta-analysis remain essential for synthesizing existing research on experimental vaccination/challenge trials. By examining data across studies, it becomes possible to identify which variables consistently drive protection, a question that individual trials cannot answer.

To address this knowledge gap, this systematic review and meta–analysis provide a quantitative synthesis of the available data on *Flavobacterium* vaccines in teleost fish. We evaluate how study characteristics, including pathogen species, vaccine type, and route of administration, influence protective efficacy, identify factors associated with greater protection, and provide evidence–based recommendations for future vaccine development and application.

## 2 Methods

### 2.1 Study design

This study was performed according to the Cochrane Handbook for Systematic Reviews of Interventions ^57^ and adheres to the structured and reporting guidelines outlined in the Preferred Reporting Items for Systematic Reviews and Meta-Analyses (PRISMA) 2020 ^58^.

### 2.2. Search strategy and information sources

To ensure a comprehensive retrieval of relevant literature, a systematic search was performed across three primary bibliographic databases: PubMed, Web of Science (All databases), and Scopus. The search covered the period from January 1, 2000, to March 31, 2026. The search string utilized a combination of Boolean operators (AND/OR) and MeSH terms (where applicable) focusing on four core components: (1) target pathogen (e.g., *Flavobacterium columnare*, *F. psychrophilum*); (2) host (e.g., fish, teleost, aquaculture, salmonids, tilapia); (3) intervention (e.g., vaccine, immunisation, bacterin, recombinant, DNA vaccine); and (4) outcome (e.g., efficacy, relative percent survival, mortality, protection). A detailed summary of the search strings for each database is provided in **Supplementary File S1**.

### 2.3. Eligibility criteria and study selection

Records published between January 2000 and March 2026 were considered. All retrieved records were imported into EndNote X9 for automated and manual deduplication. Two independent reviewers screened titles and abstracts against predefined eligibility criteria, followed by full-text assessment of potentially eligible studies. Disagreements were resolved through discussion or consultation with a third senior reviewer.

Studies were included if they: (1) were original, peer-reviewed experimental trials evaluating vaccines against *Flavobacterium* species in fish; (2) involved any fish species susceptible to *Flavobacterium* infection; (3) assessed administration of a vaccine regardless of type or route; (4) included a concurrent unvaccinated or sham-vaccinated control group subjected to the same challenge conditions; and (5) reported survival or mortality data sufficient to calculate RPS or risk ratio (RR). Only studies with full texts available in English were included. However, review articles, conference abstracts, studies lacking a formal challenge model, and trials involving co-infections were excluded.

### 2.4 Data extraction and management

Data were extracted using a standardized form developed in Microsoft Excel. The following information was recorded: (1) study characteristics, including first author, publication year, and country; (2) fish species; (3) *Flavobacterium* species and strain; (4) vaccination details, including vaccine type, administration route, and use of the boosters or adjuvants; (5) outcome measures, including numbers of dead and total fish in vaccinated and control groups, and RPS values.

### 2.5. Risk of bias assessment

The risk of bias of the included studies was assessed using the Cochrane Risk of Bias 2 (RoB 2) tool ^59^, adapted for experimental fish vaccine trials. The following domains were evaluated: (1) randomization process; (2) missing outcome data; (3) outcome measurement; and (4) selection of reported results. Each domain was rated as “low risk of bias”, “some concerns”, or “high risk of bias”.

### 2.6 Effect size calculation

The primary effect size was the RR. For descriptive interpretation, RPS was computed for each trial as ^49^:

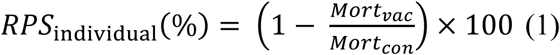

where *Mortvac* and *Mortcon* are the mortality rates in the vaccinated and control groups, respectively, calculated as *Mort_vac_* = *d_v_*/*n_v_* and *Mort_con_* = *d_c_*/*n_c_*, where *d_v_* and *n_v_* are the number of dead and total fish in the vaccinated group, and *d_c_* and *n_c_* are the number of dead and total fish in the control group.

For each study, the numbers of dead and total fish in the vaccinated and control groups were extracted. To accommodate zero-cell counts, a continuity correction of 0.5 was added to every cell of the 2×2 table ^60–62^. Because both the dead and surviving cells in each group receive 0.5, the total number of fish increases by 1. For the calculation of the RR, the correction was implemented as:

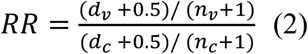

The standard error of the log risk ratio was then calculated as ^61^:

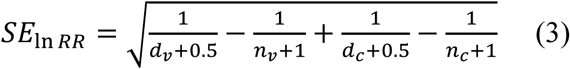

The 95% confidence interval (CI) for the RR was obtained by exponentiating the log-transformed limits ^60^:

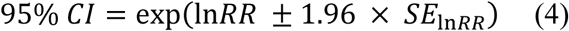

When presenting pooled RPS, the pooled RPS was derived directly from the pooled RR obtained from the random-effects meta-analysis:

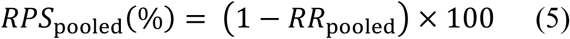

When the number of deaths was not stated directly, we obtained it in one of two ways. If the paper reported RPS together with the control-group mortality or survival rate, the number of deaths in the vaccinated group (*d_v_*) was back-calculated as:

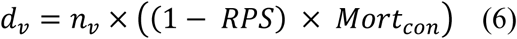

If survival rate was reported instead, *Mort_con_* = 1-survival rate. Values were rounded to the nearest integer when necessary. If mortality or survival data were shown only in figures (e.g., survival curves, bar charts), we extracted the endpoint mortality or survival rate with WebPlotDigitizer (version 4.8).

All statistical inferences (pooled estimates, heterogeneity, subgroup differences, publication bias) were based on RR.

### 2.7 Data Analysis

All analyses were conducted in R using the meta packages (version 4.3.3, R Core Team, Vienna, Austria), in accordance with the PRISMA guidelines. Meta-analyses were performed using random-effects models with the DerSimonian–Laird estimator for the between-study variance (*τ^2^*). Effect sizes are reported as RRs with 95% CIs. Heterogeneity was assessed using Cochran’s *Q* test, Higgins’ *I*^2^ statistic, and *τ^2^*. An *I*^2^ value greater than 50% was considered indicative of substantial heterogeneity. Galbraith (radial) plots were used to identify potential outliers and influential studies.

Subgroup analyses were conducted to evaluate the effects of challenge species, vaccine type, administration route, booster vaccination and adjuvant use on vaccine efficacy. Subgroup differences were tested using Cochran’s *Q* test. For pathogen–host combinations with at least two independent trials, separate random-effects meta-analyses were performed within each combination to obtain pooled RRs, which are visualised in a heatmap. For studies with multiple treatment arms sharing a common control group, each arm was treated as an independent comparison; this approach avoids overcounting the control group and follows standard practice for such designs.

Publication bias was assessed by visual inspection of funnel plots and Egger’s regression test ^63^. Sensitivity analyses were conducted to examine the robustness of the findings, including leave-one-out analysis (sequential exclusion of each study), exclusion of studies with 100% control-group mortality, and exclusion of small-sample studies (any group with fewer than 50 fish). All sensitivity analyses used the same random-effects model as the primary analysis. All statistical tests were two-sided, and a *P* value <0.05 was considered statistically significant unless otherwise specified.

## 3 Results

### 3.1. Study selection and characteristics

The study selection process is summarized in **Figure 1**. A total of 1,044 records were identified through database searches, including PubMed (n=159), Web of Science (n=596), and Scopus (n=289). After removal of 421 duplicate records, 623 records remained for title and abstract screening. Following screening and eligibility assessment, 60 studies were included in the systematic review, of which 59 met the criteria for meta-analysis (one of which was partially included ^39^), representing 377 independent vaccine challenge comparisons (**Figure 1**).

**Figure 1.**
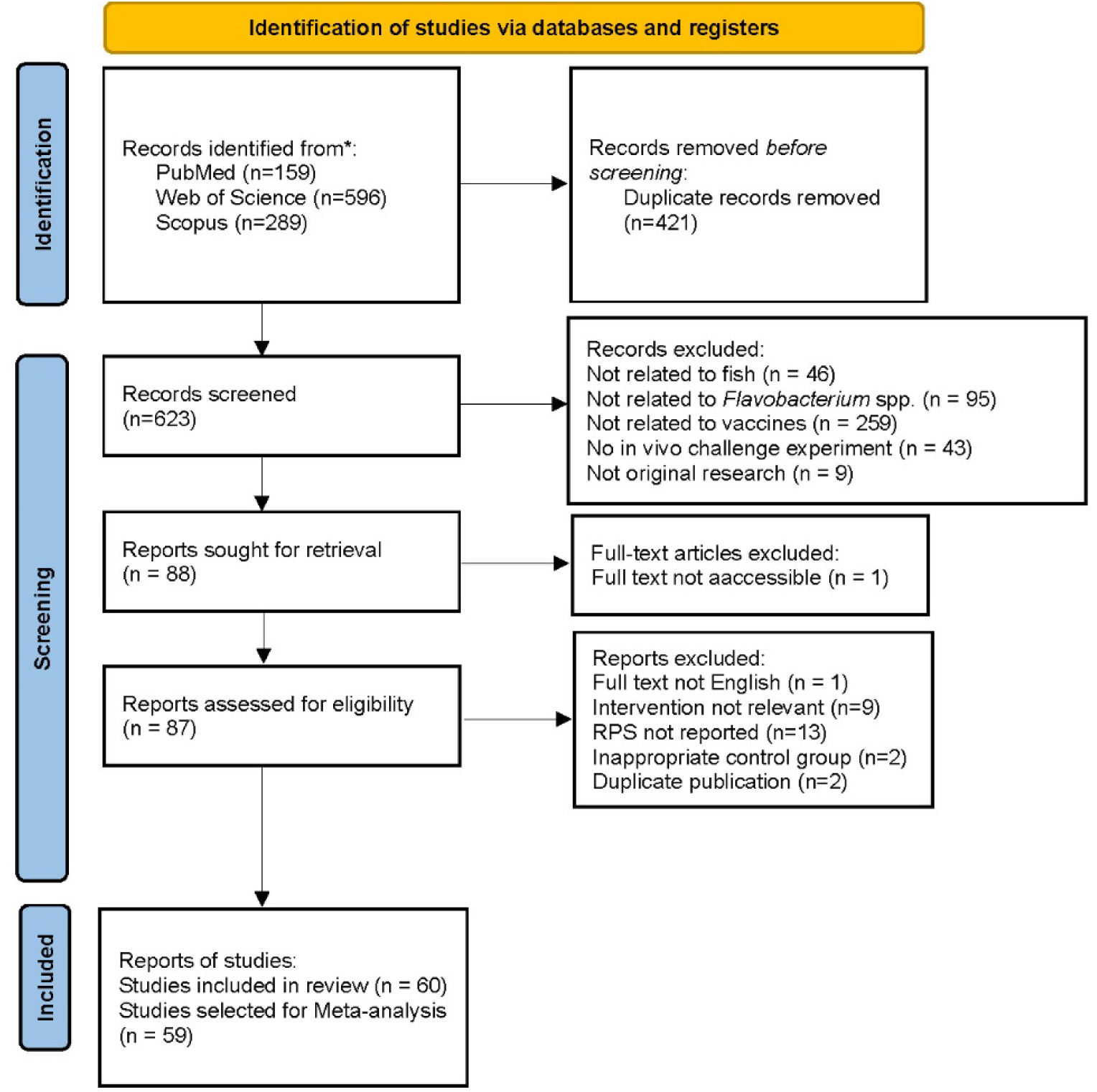
PRISMA flowchart for included studies. Of the 59 studies included in the meta–analysis, 58 met the full criteria for quantitative synthesis, and one study was included only for particular comparisons (not all comparisons from that study were used).

Detailed characteristics of the included studies are provided in **Table S1**. The studies encompassed multiple fish hosts, *Flavobacterium* species, vaccine platforms, routes of administration, booster vaccination and adjuvant use. Three studies could not be confidently assigned to a specific *Flavobacterium* species based on current taxonomic classification and was therefore included only in the overall pooled RR analysis, but excluded from pathogen-specific subgroup analyses.

### 3.2. Heterogeneity and publication bias

The distribution of individual, unweighted RPS values is presented in **Figure 2A**. Across all included comparisons, the median RPS was 41.7% (interquartile range [IQR]: 15.4–66.0%), with values ranging from −40% to 99.1%. A small number of comparisons reported negative RPS values, indicating higher mortality in vaccinated fish than in corresponding unvaccinated groups. Substantial between-study heterogeneity was detected in the overall meta-analysis (*I*^2^ = 89.5%, *τ*^2^ = 0.125; **Figure 5**), indicating marked variation in vaccine efficacy across trials. The random-effects Galbraith plot (**Figure 2B**) showed broad dispersion, with many studies lying outside the 95% confidence limits, consistent with the high *I^2^*value.

**Figure 2.**
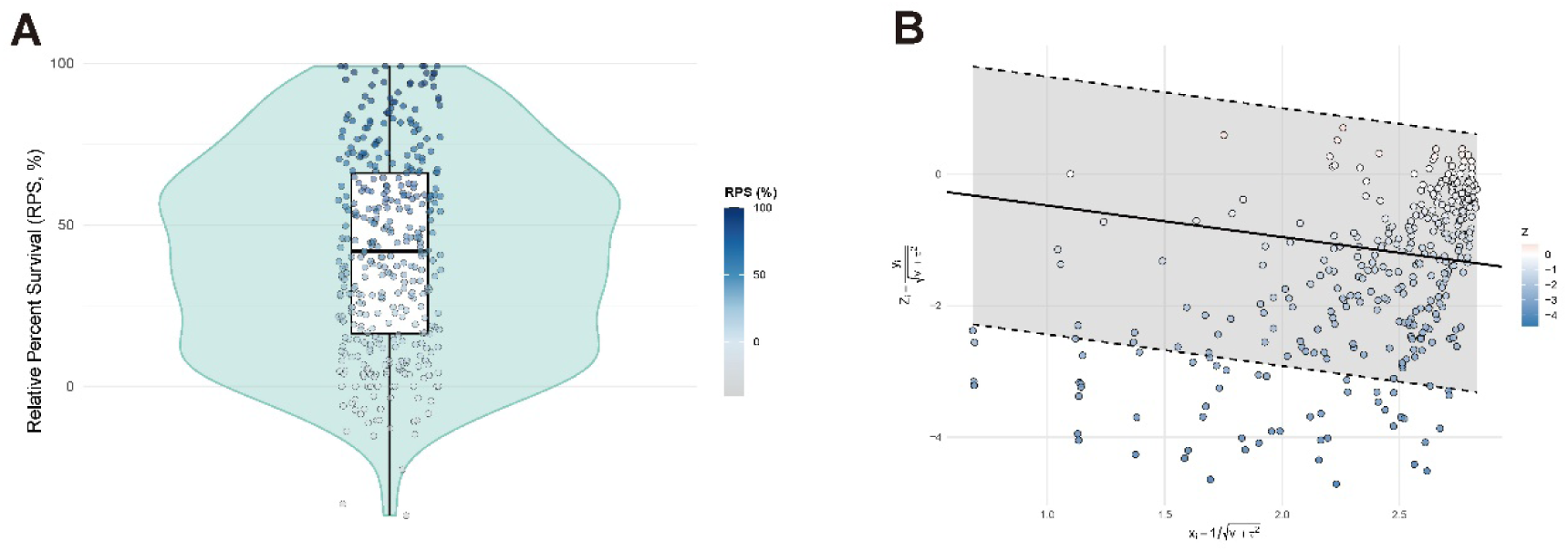
(A) Violin plot showing the distribution of relative percent survival (RPS) across all included trials (n = 377). The black bold line within the box indicates the median, and the box bounds represent the interquartile range. The color of jittered points reflects the RPS value, ranging from gray (low RPS) to dark blue (high RPS) as shown in the color gradient bar on the right. **(B)** Galbraith plot for heterogeneity assessment.

Publication bias was evaluated by the funnel plot (**Figure S1**) which showed asymmetry, and Egger’s regression test which was significant (t = −12.0, df = 375, *P* < 0.001), indicating potential publication bias or small-study effects.

### 3.3. Quality assessment and risk of bias

The risk of bias summary for the 60 included studies is presented in **Figure 3**. No study was judged as high risk. Randomization procedures were rated as low risk in 43.3% of studies and some concerns in 56.7%. Missing outcome data was low risk in 98.3% and some concerns in 1.7%. Outcome measurement was low risk in all studies (100%). Selection of the reported result was low risk in 96.7% and some concerns in 3.3%. Detailed risk of bias judgements for each individual study are provided in **Figure S2**.

**Figure 3.**
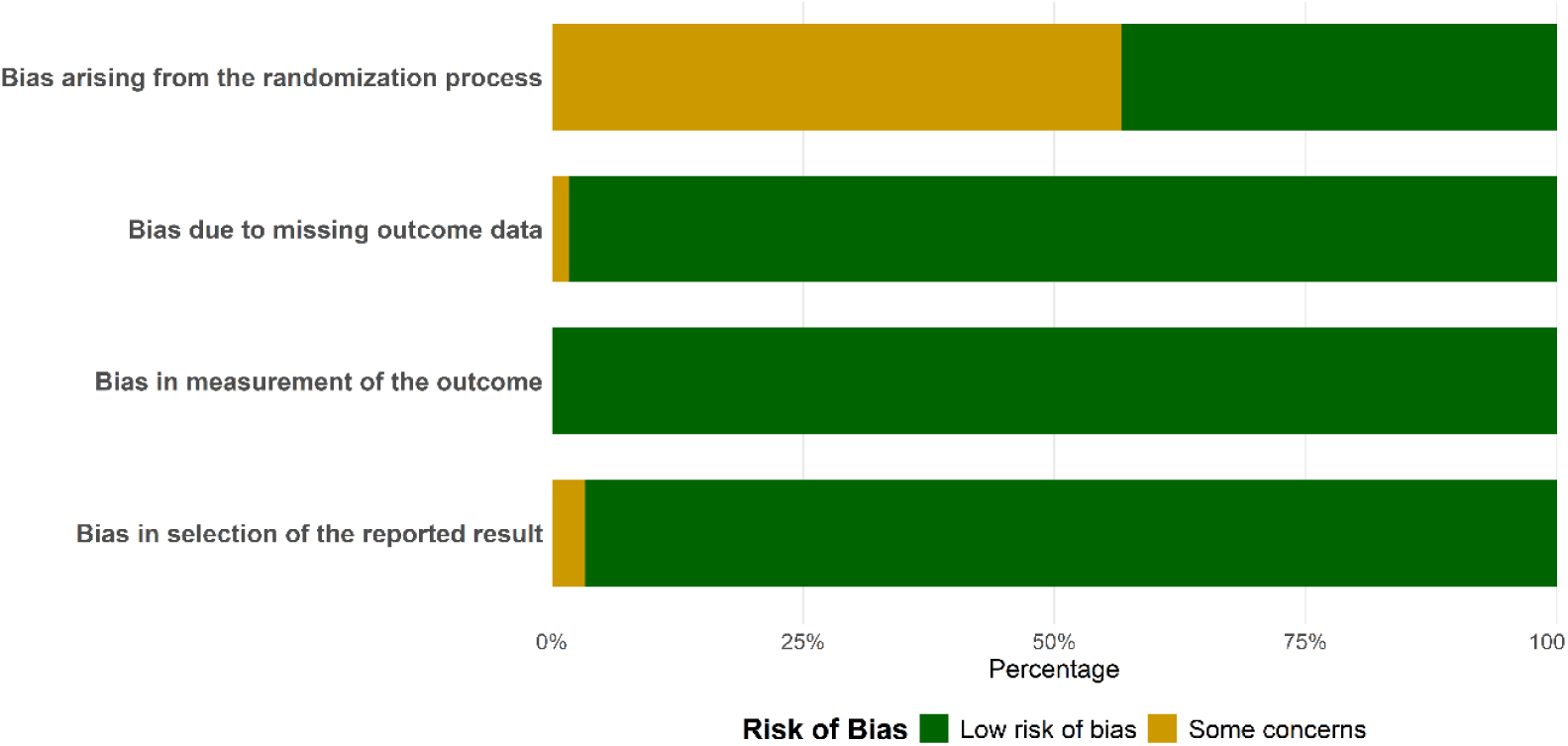
Risk of bias summary for the 60 included studies.

### 3.4. Meta-analysis and subgroup analysis

#### 3.4.1. Host-pathogen specificity

Pooled vaccine efficacy across pathogen–fish host combinations is presented in **Figure 4**, showing marked variation in RPS between groups. For *F. psychrophilum*, RPS ranged from 44.7% in rainbow trout to 82.6% in salmon. Among CCB species, *F. davisii* consistently showed strong protection across hosts (RPS >80%). *F. columnare* was evaluated only in grass carp (30.6%). *F. covae* showed the highest protection in tilapia (RPS = 59.5%), followed by channel catfish (41.9%) and grass carp (31.5%), but performed poorly in Asian seabass (29.7%) and zebrafish (18.3%). However, it should be noted that the tilapia estimate was based on only two comparisons. *F. oreochromis* showed similar RPS in tilapia and Asian seabass (36.5% vs. 34.9%). *F. johnsoniae* in grass carp yielded 43.7%, though all comparisons were from a single study ^64^.

**Figure 4.**
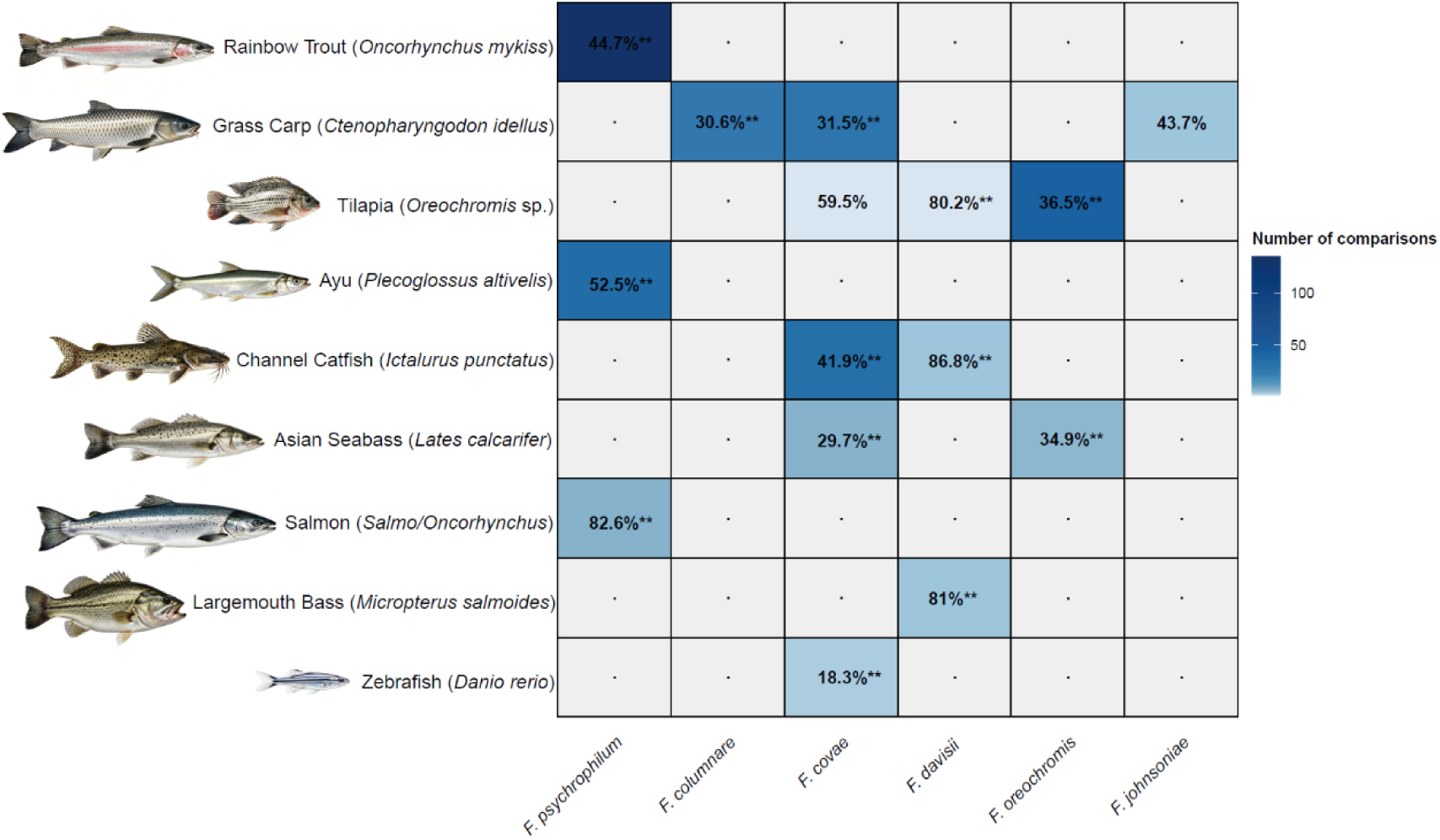
Heatmap of pooled vaccine efficacy for pathogen–fish host combinations with at least two independent comparisons. Each cell represents the pooled RPS for a given *Flavobacterium* species and fish host. The colour intensity represents the number of comparisons (k) contributing to each pooled estimate: darker blue indicates a larger number of comparisons. Gray cells indicate no data or *k*=1 (e.g., *F. araucananum* ^90^). Asterisks indicate the significance of the pooled RPS: ** *p*<0.01, * *p*<0.05.

**Figure 5.**
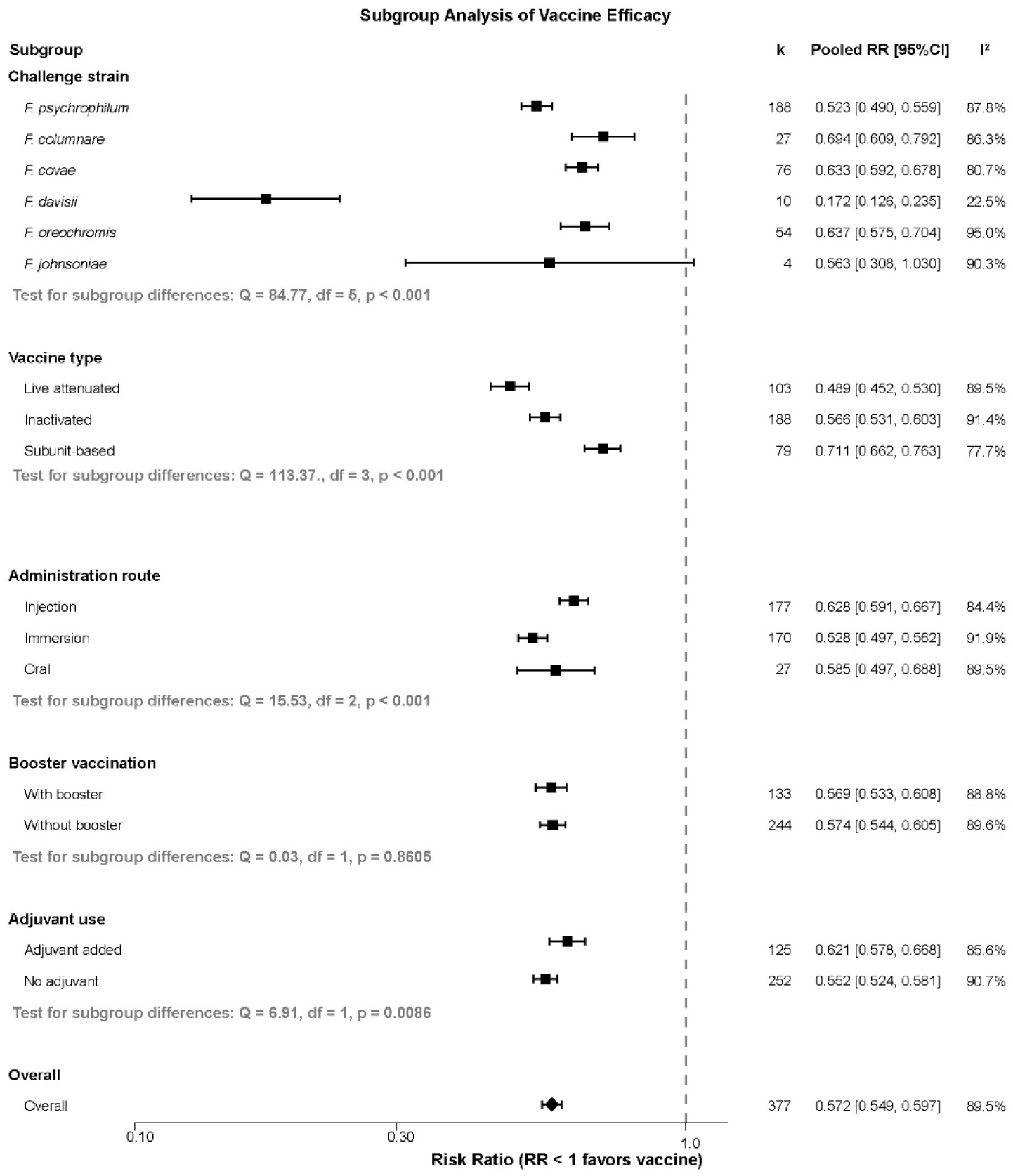
Subgroup analysis of vaccine efficacy by challenge strain, vaccine type, administration route, booster vaccination and adjuvant use. Vaccine type subgroup analyses were limited to live attenuated, inactivated, and subunit vaccines. Only comparisons with a single vaccine type and single route were included; combination/multicomponent vaccines and mixed routes were excluded. Booster was defined as a second immunisation ≥1 week after the primary dose; adjuvant referred to explicitly added immunostimulants (e.g., mineral oil, Freund’s complete/incomplete adjuvant).

#### 3.4.2. Subgroup analysis

Pooled efficacy estimates stratified by pathogen species, vaccine type, administration route, booster vaccination and adjuvant use are shown in **Figure 5**. The pooled RR varied significantly among *Flavobacterium* species (test for subgroup differences: Q = 84.77, *P* <0.001), ranging from 0.172 (95% CI: 0.126–0.235; RPS = 82.8%) for *F. davisii* to 0.694 (95% CI: 0.609–0.792; RPS = 30.6%) for *F. columnare*. The overall RR was 0.572 (95% CI: 0.549–0.597; RPS = 42.8%). Other species showed RR: *F. psychrophilum* 0.523 (0.490–0.559; RPS = 47.7%), *F. covae* 0.633 (0.592– 0.678; RPS = 36.7%), *F. oreochromis* 0.637 (0.575–0.704; RPS = 36.3%) and *F. johnsoniae* 0.563 (0.308–1.030; RPS = 43.7%). Heterogeneity was substantial in all subgroups except *F. davisii* (*I*² = 22.5%).

For vaccine type, a significant subgroup difference was observed (Q = 113.37, *P* <0.001). Live attenuated vaccines showed the lowest RR with highest protective efficacy (0.489, 95% CI: 0.452–0.530; RPS = 51.1%), followed by inactivated vaccines (0.566, 95% CI: 0.531–0.603; RPS = 43.4%) and subunit vaccines (0.711, 95% CI: 0.662–0.763; RPS = 28.9%). Heterogeneity was substantial in all subgroups.

For the administration route, a significant subgroup difference was observed (Q = 15.53, *P* <0.001). Injection resulted in an RR of 0.628 (95% CI: 0.591–0.667; RPS = 37.2%), immersion yielded an RR of 0.528 (95% CI: 0.497–0.562; RPS = 47.2%), and oral administration showed an RR of 0.585 (95% CI: 0.497–0.688; RPS = 41.5%). Heterogeneity remained high across all routes (*I*² = 84.4%, 91.9% and 89.5%, respectively).

For booster vaccination, no significant difference was detected between vaccination with and without a booster dose (Q = 0.03, df = 1, *P* = 0.8605). The pooled RR was 0.569 (95% CI: 0.533– 0.608; RPS = 43.1%) for the booster group and 0.574 (95% CI: 0.544–0.605; RPS = 42.6%) for the without booster group.

For adjuvant use, a significant subgroup difference was observed (Q = 6.91, *P* = 0.0086). Vaccines formulated with adjuvant yielded an RR of 0.621 (95% CI: 0.578–0.668; RPS = 37.9%), compared with 0.552 (95% CI: 0.524–0.581; RPS = 44.8%) for those without adjuvant. Heterogeneity was similarly high in both categories (*I*² = 85.6% and 90.7%, respectively).

#### 3.4.3. Detailed comparison-level analysis of individual pathogen species

For *F. psychrophilum*, detailed comparison-level results stratified by fish host (rainbow trout, salmon, and ayu) are provided in **Supplementary Table S2**. For rainbow trout (k = 145), individual RRs ranged from 0.01 to 1.36; for salmon (k = 7), from 0.03 to 0.54; and for ayu (k = 36), from 0.09 to 1.00. The majority of comparisons favoured vaccination (RR <1), with weights broadly distributed.

Detailed forest plots showing the contribution of each individual comparisons are presented for *F. columnare* (**Figure 6**), *F. covae* (**Figure 7**), *F. davisii* (**Figure 8**), and *F. oreochromis* (**Figure 9**). For *F. columnare*, all 27 trials were conducted in grass carp (*Ctenopharyngodon idella*). The RRs of individual trials ranged from 0.06 to 1.11, with most studies favouring vaccination (RR <1). For *F. covae*, which had the largest number of trials (k = 76), RRs ranged from 0.19 to 1.04. For *F. davisii*, all 10 trials used live attenuated vaccines administered by immersion, and all showed strong protection (RR 0.05–0.28). For *F. oreochromis*, all 54 trials employed inactivated vaccines, with RRs ranging from 0.18 to 1.11. *F. johnsoniae* (k = 4) is not shown separately as all trials were from a single study^64^; its pooled estimate is presented in **Figure 5**.

**Figure 6.**
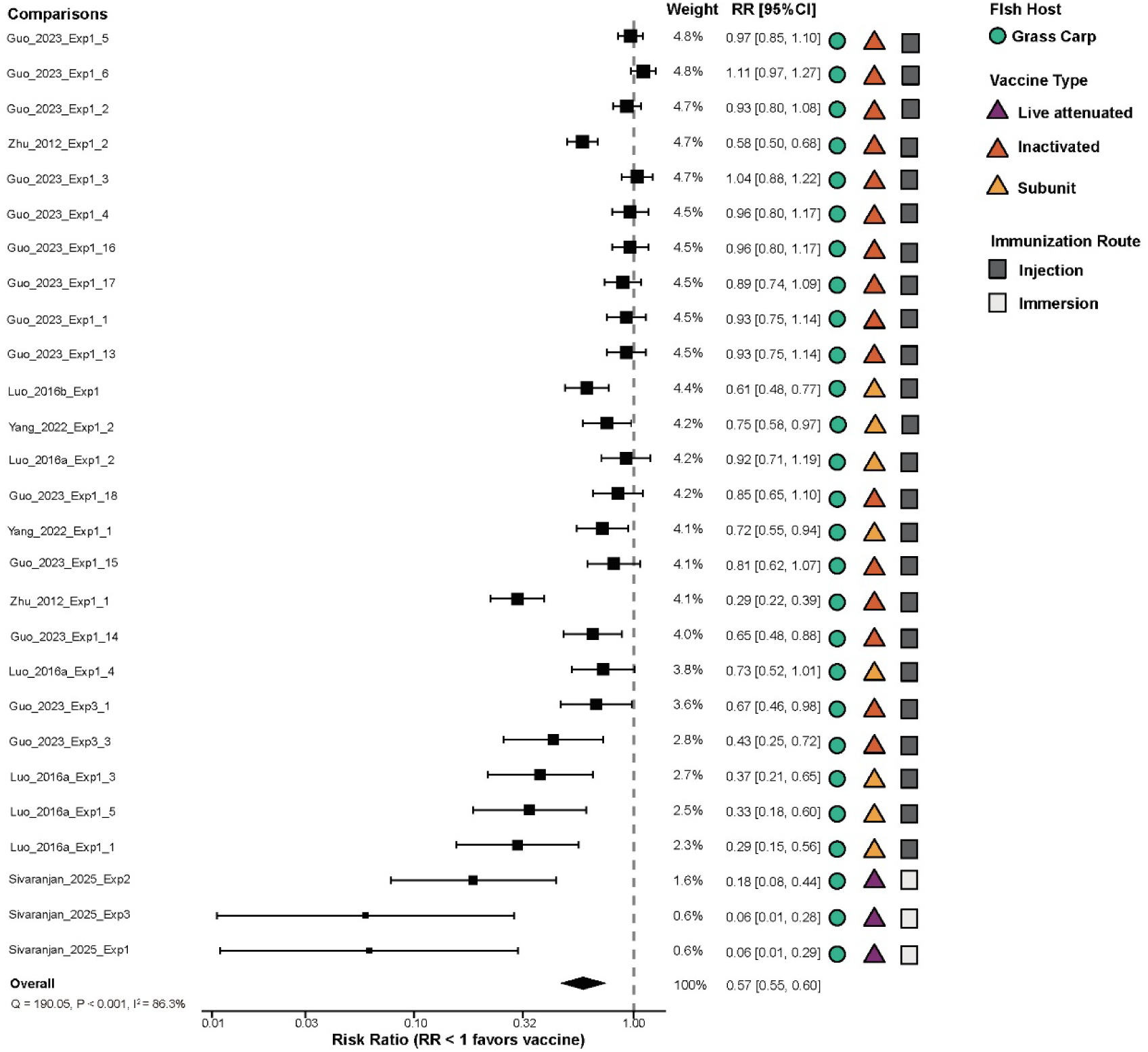
Forest plot of pooled vaccine efficacy against *F. columnare* in grass carp, stratified by vaccine type and route of administration.

**Figure 7.**
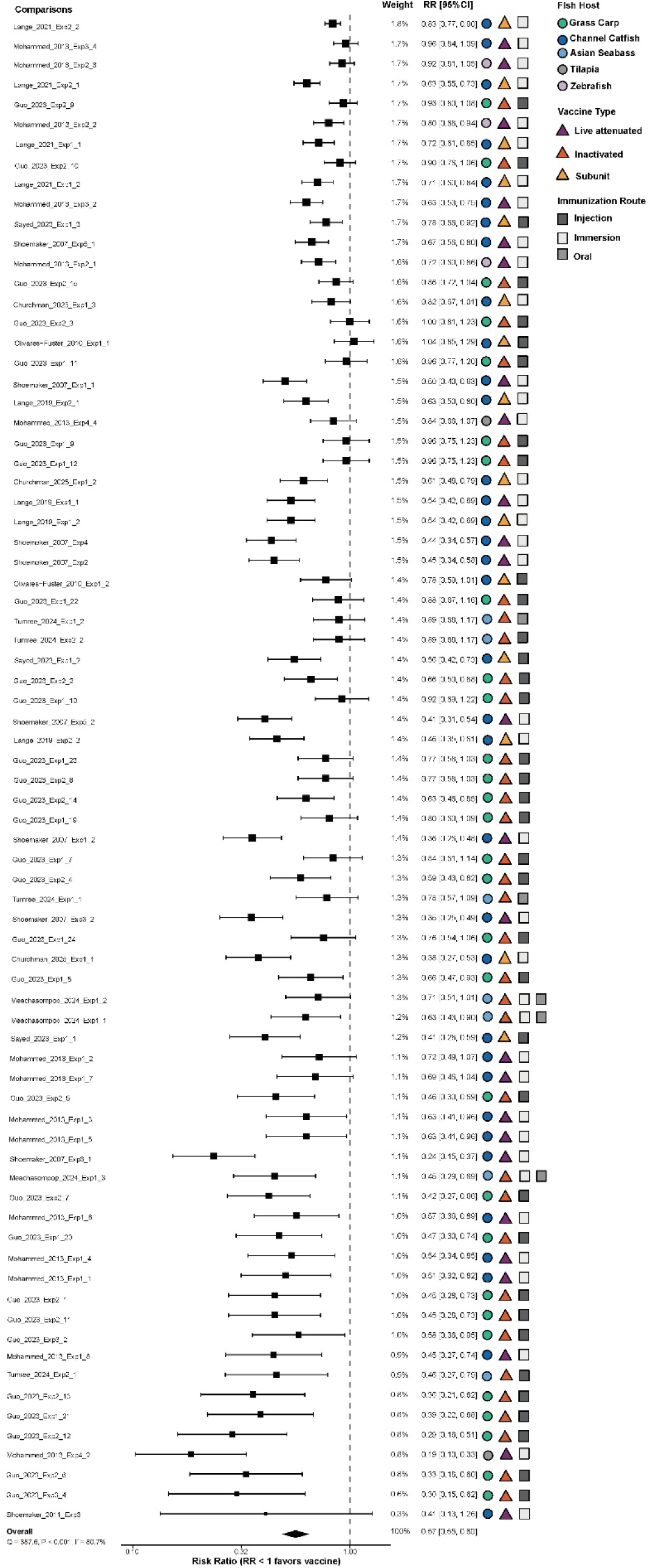
Forest plot of pooled vaccine efficacy against *F. covae*, stratified by fish host, vaccine type, and route of administration.

**Figure 8.**
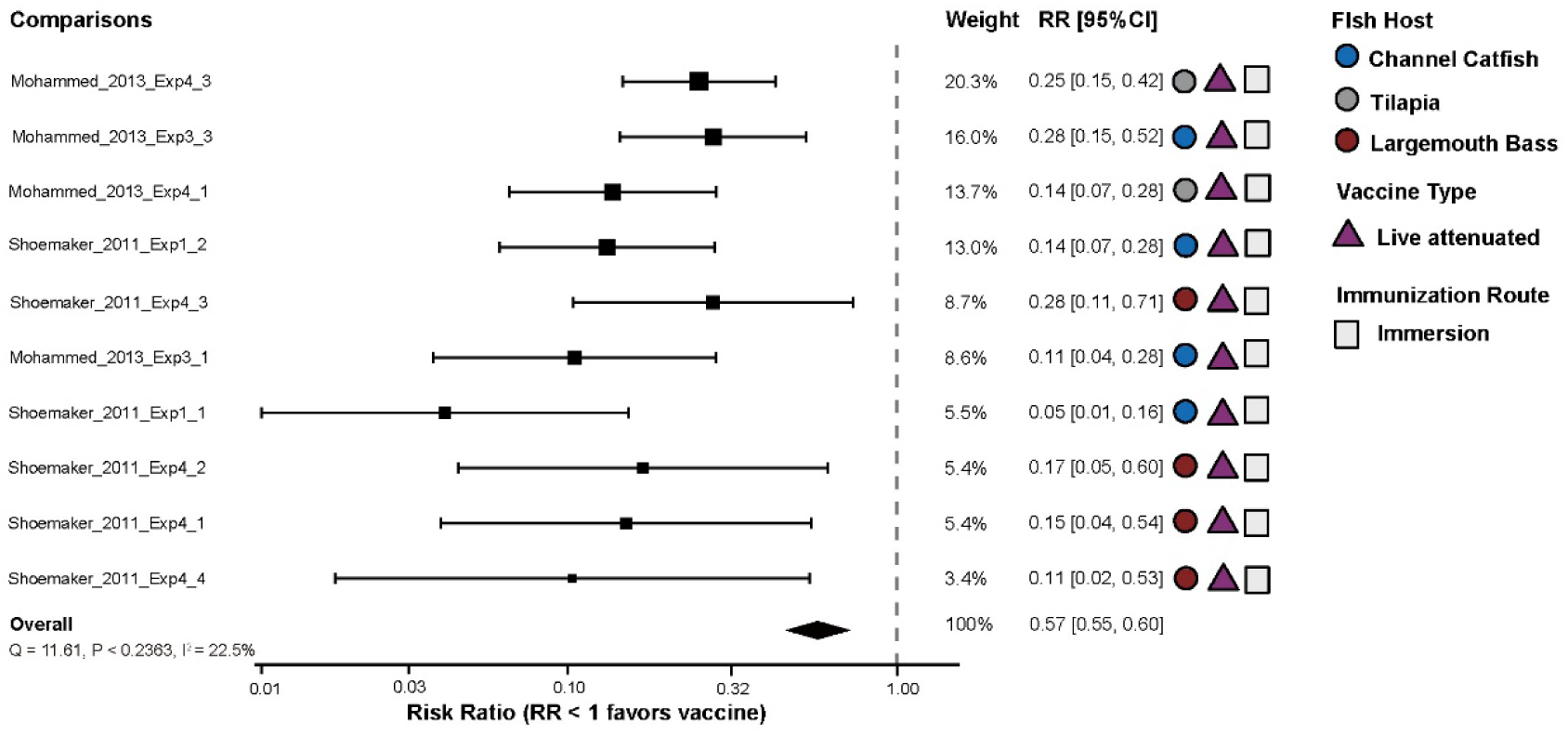
Forest plot of pooled vaccine efficacy against *F. davisii*, stratified by fish host.

**Figure 9.**
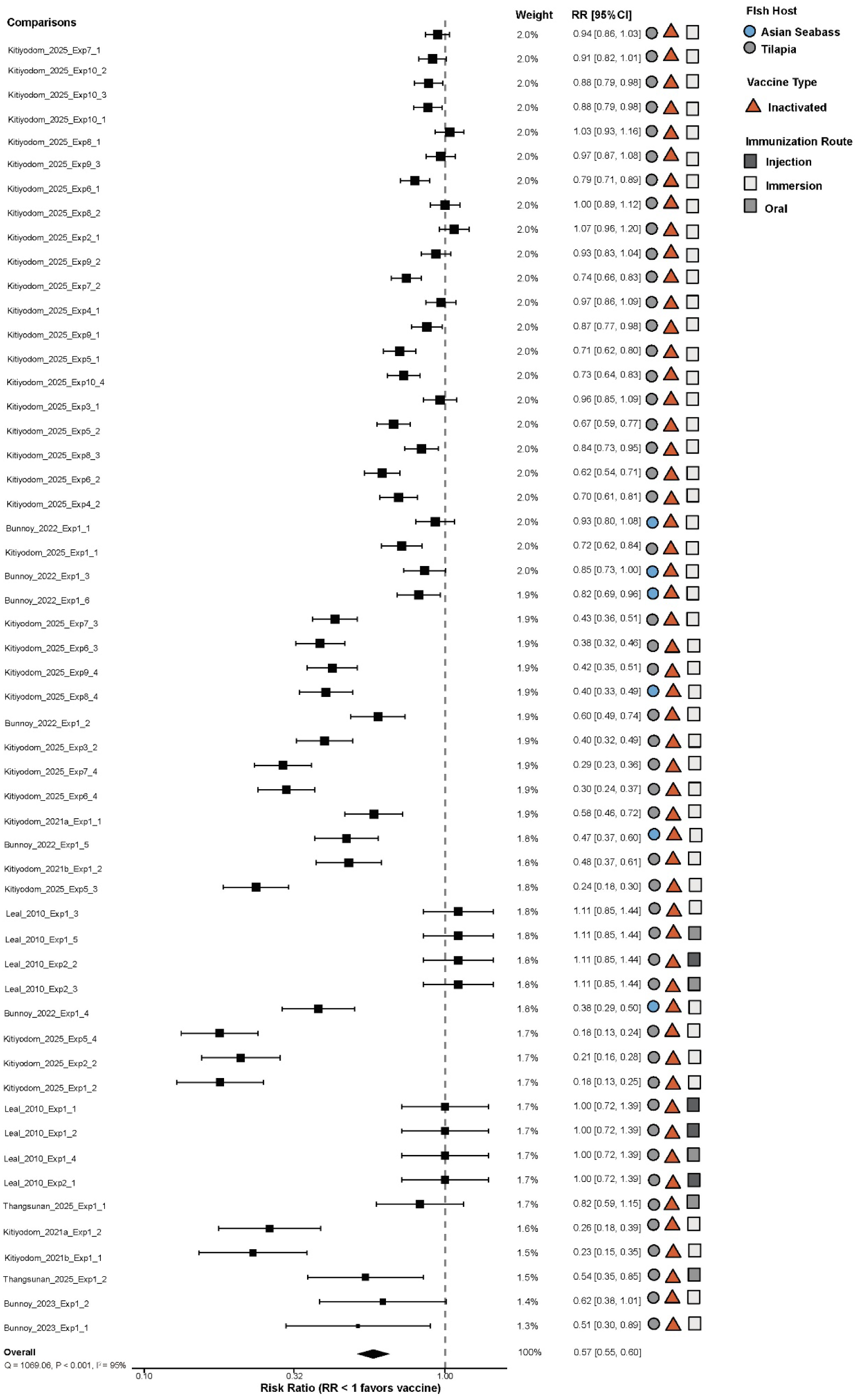
Forest plot of pooled vaccine efficacy against *F. oreochromis*, stratified by fish host and route of administration.

### 3.5. Sensitivity analysis

Sensitivity analyses confirmed the robustness of the primary estimate (**Figure 10**), with all sensitivity analyses yielding pooled RRs consistent with this overall estimate (range: 0.54– 0.57). Excluding studies with 100% control mortality (k = 11) produced a pooled RR of 0.56 (95% CI: 0.54–0.59; *I*^2^ = 89.4%), while excluding small-sample comparisons (n <50 per group, k = 143) resulted in a pooled RR of 0.54 (95% CI: 0.51–0.56; *I*^2^ = 92.0%). Leave-one-out analysis yielded a narrow range of pooled RRs (0.55–0.59) around the overall RR (**Figure S3**), with *I*² values remaining similarly high (range: 87.2%–90.1%).

**Figure 10.**
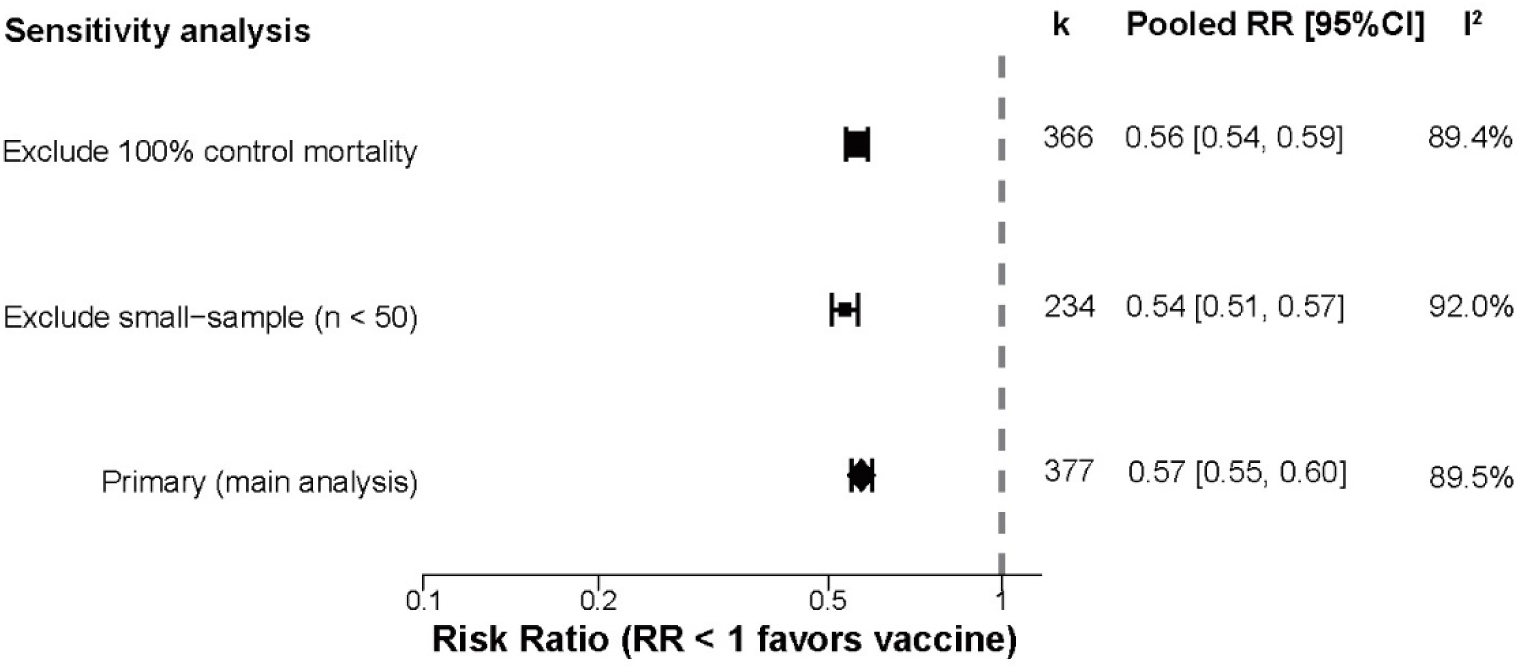
Sensitivity analysis result. The pooled RR remained stable after excluding studies with 100% control mortality, excluding small-sample comparisons (n < 50 per group).

### 3.6. Cross-protection

A total of 24 trials from four studies evaluated cross-species protection (**Figure S4, Table S3**) ^38,39,51,65^. Vaccines derived from *F. columnare* and challenged against *F. covae* produced RPS values ranging from 4.4% to 76.8% (**Figure 10**). The licensed live-attenuated vaccine AQUAVAC COL achieved high protection (RPS = 62.5%) in channel catfish. In contrast, the attenuated mutant FCRR, the active ingredient in this vaccine, showed lower efficacy, with RPS values of 4.4% in channel catfish, 32% in zebrafish, and 16.1% in tilapia. An inactivated *F. columnare* bacterin provided 42% RPS against *F. covae* in grass carp.

The reverse combination, *F. covae*-based vaccines against *F. columnare*, was evaluated in a single comparison and gave an RPS of 55% in grass carp, which was markedly higher than the homologous protection of *F. columnare* (RPS = 32%) but still lower than the homologous protection of *F. covae* (RPS = 71%).

Cross-protection against *F. davisii* was also examined. Vaccines derived from *F. columnare* and challenged with *F. davisii* produced RPS values between 72.7% and 96.4%. The FCRR vaccine achieved RPS of 72.7% in channel catfish and 75.5% in tilapia, whereas AQUAVAC-COL gave RPS of 74.6–92.6% in largemouth bass across four dose levels. Vaccines based on *F. covae* against *F. davisii* yielded RPS of 86.9% in channel catfish and 90.9% in tilapia, indicating strong cross-protection.

### 3.7. Polyvalent vaccine efficacy

Twenty experimental comparisons from eight studies evaluated polyvalent or bivalent vaccines containing *Flavobacterium* antigens (**Table S3**). For *F. psychrophilum*, a total of 11 experimental comparisons evaluated polyvalent and bivalent vaccines. In rainbow trout, inactivated vaccines gave RPS values of 25%–100% for multivalent formulations ^48,66,67^ and 79.5% for a bivalent vaccine ^68^. In Atlantic salmon, the multivalent vaccine yielded RPS values of 75%– 95% against a homologous strain ^69^.

For the bivalent vaccines against *F. covae*, RPS values ranged from 11.3% to 59.7%. In channel catfish, live-attenuated bivalent vaccines targeting *F. covae* and *E. ictaluri* achieved RPS values of 33%–59.7%, compared with 50%–76.8% for corresponding monovalent *F. covae* vaccines. In Asian seabass, an inactivated bivalent vaccine (containing *S. iniae* and *F. covae*) administered by immersion and oral gave RPS values of 28.41%–55.26% (no monovalent control). Another trial in Asian seabass compared monovalent and bivalent inactivated *F. covae* vaccines via oral or injection routes: the monovalent vaccine produced RPS of 23.1%–54.4%, while the bivalent formulation yielded 11.3% RPS by both routes. For *F. oreochromis*, a bivalent nanovaccine targeting *F. oreochromis* and *F. orientalis* achieved 38.89% RPS, compared with 50% for the monovalent *F. oreochromis* vaccine.

### 3.8. Intrinsic virulence of *Flavobacterium* species

The distribution of control-group mortality, used as an indicator of pathogen virulence in the absence of vaccination, is presented in **Figure 11**. Median mortality differed among *Flavobacterium* species, ranging from 43.8% (IQR: 36.7–43.8%) for *F. davisii* to 90.0% (IQR: 83.8–93.9%) for *F. columnare*. *F. covae* and *F. johnsoniae* showed similarly high medians (83.3% and 85.0%, respectively). Intermediate values were observed for *F. psychrophilum* (77.3%, IQR: 55.9–92%) and *F. oreochromis* (73.3%, IQR: 69–77.7%).

**Figure 11.**
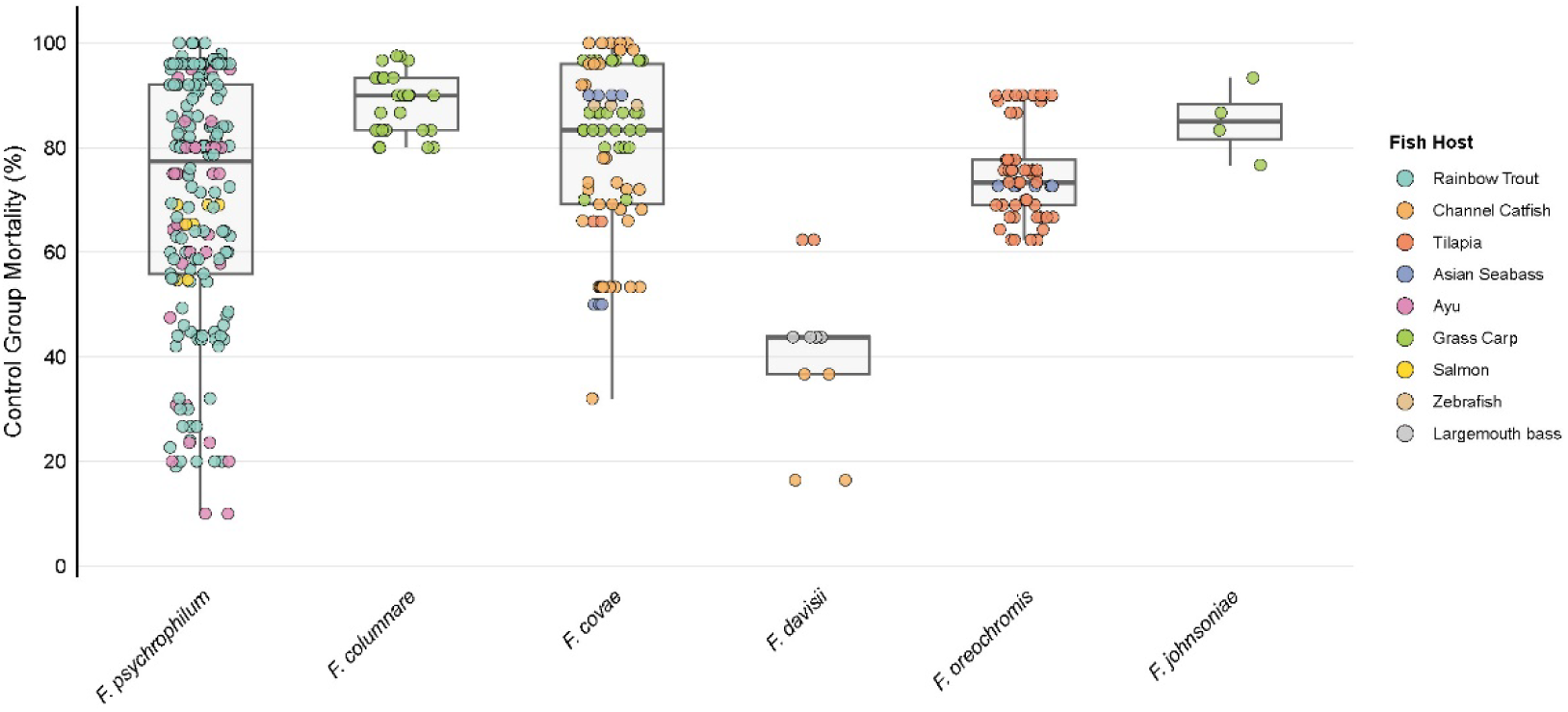
Intrinsic virulence of *Flavobacterium* species as assessed by control group mortality. The boxplots display the distribution of control mortality (%) for each pathogen species. Jittered points represent individual comparisons, colored by fish host.

## 4 Discussion

This systematic review and meta-analysis integrate vaccine efficacy data against *Flavobacterium* infections in fish, yielding an overall pooled RPS of 42.8%. This finding confirms that vaccination is broadly effective in reducing mortality risk caused by *Flavobacterium* species, with coverage extending beyond individual trials or previous narrative reviews. Previous narrative reviews have summarized available vaccine technologies and reported protection rates from individual studies, but have not quantitatively synthesized these data ^70–72^. Individual trials are typically constrained by limited sample sizes and are therefore subject to greater random variation. Moreover, a single trial usually evaluates one vaccine product against a specific pathogen, in a particular host ^73–75^, making its results difficult to project directly to other conditions and compare the relative merits of different vaccine products. By pooling data from 377 independent comparisons, this study not only provides robust evidence to date for the protective efficacy of vaccines against *Flavobacterium* species, but also systematically evaluates multiple factors that may affect protection. However, the substantial heterogeneity underlying the pooled effect (*I*² = 89.5%) indicates that the overall estimate is a weighted average that masks considerable variability and that the true efficacy of vaccination is highly dependent on the specific conditions under which the vaccine is applied.

Protective efficacy differed markedly among *Flavobacterium* species. *F. davisii* showed the highest protection (pooled RPS = 82.8%), followed by *F. psychrophilum* (47.7%) and *F. johnsoniae* (43.7%). In contrast, *F. columnare* (30.6%), *F. covae* (36.7%), and *F. oreochromis* (36.3%) exhibited relatively lower protection. Notably, heterogeneity remained high even within individual species, which may partly reflect differences in host specificity and antigenic diversity. *F. psychrophilum* (*k* = 188) was the largest subgroup and also exhibited significant heterogeneity (*I*² = 87.8%). This high heterogeneity is consistent with the documented host specificity of *F. psychrophilum*: strains isolated from rainbow trout are nearly avirulent for ayu, and vice versa, and this host-specific virulence is closely linked to O-antigen serotype and clonal complex ^76,77^. Multiple serotypes exist in *F. psychrophilum* based on O-antigen polysaccharide structure, and the degree of cross-protection between serotypes varies ^48,67,68^. Therefore, vaccines formulated from a single serotype strain may not provide the same level of protection against all circulating strains, indicating that the inclusion of multiple serotypes in vaccine formulations deserves further investigation. Among the CCB group, *F. covae* accounted for the largest number of comparisons (*k* = 76), predominantly in channel catfish. Its pooled RR was 0.633 (95% CI: 0.592–0.678), with high subgroup heterogeneity (*I*² = 80.7%), indicating substantial variability in protection that is also influenced by host specificity. The RPS of *F. covae* vaccines reached 59.5% in Tilapia (*Oreochromis* sp.), whereas it only yielded 18.3% in zebrafish (*Danio rerio*). This host-related variability in vaccine efficacy suggests that vaccines intended for a given host are best developed using strains isolated from that same host, while cross-host protection will likely depend on identifying conserved antigens shared across serotypes. *F. davisii* exhibited the lowest pooled RR among the CCB group, with the lowest subgroup heterogeneity (*I*² = 22.5%) and the most stable protection; however, the number of trials was limited (*k* = 10), and the results may be affected by chance or by underreporting of trials with poor efficacy. Similarly, evidence for *F. johnsoniae* and *F. araucananum* remains scarce, and further trials may be required to evaluate the efficacy of the vaccine against these species. *F. columnare* and *F. oreochromis* showed relatively low protective efficacy, with *F. columnare* being the lowest (RR = 0.694, 95% CI: 0.609–0.792; *I*² = 86.3%). *F. oreochromis* also exhibited high heterogeneity (*I*² = 95.0%), indicating that the protection provided by current vaccines against this species is inconsistent, underscoring the urgent need to develop more effective and reliable vaccines. All *F. columnare* comparisons were conducted in grass carp; the substantial heterogeneity suggests that the instability of vaccine efficacy is further influenced by factors such as vaccine type and route of administration.

The incorporation of vaccine type further contributed to the observed heterogeneity. Subgroup analysis by vaccine type showed that live attenuated ranked highest in protective efficacy (RPS = 51.1%), followed by inactivated vaccines (43.4%) and subunit vaccines (28.9%). This difference may be attributable to the broader antigenic repertoire of whole-cell vaccines, whether live or inactivated, whereas subunit vaccines typically present only a limited set of antigens and require potent adjuvants to induce an adequate immune response.

Administration route and adjuvant use significantly affected vaccine efficacy (P < 0.001 and *P* = 0.0086, respectively). Each subgroup was characterised by high heterogeneity (*I*² = 84.4– 91.9%). It remains possible that the effects of route or adjuvant were masked by other between-study variations, such as differences in challenge conditions, host species, or vaccine formulation. A notable exception was *F. oreochromis*, for which injection produced RR ≥1, indicating no protective effect, suggesting that this route may be unsuitable for this pathogen. Moreover, all vaccine trials against *F. oreochromis* to date have evaluated inactivated vaccines; live attenuated and subunit vaccines remain unexplored, and alternative vaccine types or delivery strategies warrant investigation.

Even when fish host and vaccine type were held constant, different strains of the same bacterial species still yielded substantially different levels of protection. Analysis of cross-protection data revealed that AQUAVAC-COL and FCRR, both live attenuated vaccines derived from *F. columnare*, are fully matched in host and vaccine type. Still, they differed dramatically in their cross-protective efficacy against *F. covae* in channel catfish: RPS reached 62.5% for AQUAVAC-COL but only 4.4% for FCRR ^39,51^. Such a large difference between two live attenuated vaccines originating from the same *F. columnare* lineage strongly suggests that different attenuated mutants retain or lose distinct sets of protective antigens during passage, and that careful strain selection and cross-protection were also highly important ^51^. Vaccines derived from *F. covae* consistently provided stronger cross-protection against *F. columnare* and *F. davisii* than the reverse combinations. The *F. covae* 17-23 mutant achieved RPS values of 86.9% in channel catfish and 90.9% in tilapia against *F. davisii* and 55% against *F. columnare*, all exceeding the corresponding homologous protection levels. This directional asymmetry suggests that *F. covae* may express a set of more conserved cross-protective antigens. In addition, host specificity further complicates the assessment of cross-protection. Vaccines derived from *F. columnare* conferred the highest protection against *F. covae* in catfish, but only modest protection in grass carp and tilapia, suggesting that cross-protection depends not only on the antigenic similarity between vaccine and challenge strains but also on the host specificity. In summary, cross-protection reflects the combined influence of pathogen, host, and vaccine. Conserved antigens shared across the CCB group, while necessary targets for broad-spectrum vaccine design, must still prove effective in the intended target host.

The development of polyvalent vaccines has also produced contrasting results between *F. psychrophilum* and the CCB group. For *F. psychrophilum*, polyvalent inactivated vaccines maintained high levels of protection (RPS 25–100%), with efficacy comparable to their monovalent counterparts ^48,66–69^, supporting the feasibility of combining multiple antigens in a single formulation for this pathogen. However, this finding does not readily extend to CCB. Evidence for polyvalent vaccines in CCB remains limited, and the available comparisons have generally failed to show any advantage over monovalent formulations. For example, a bivalent *F. covae* and *S. iniae* vaccine produced an RPS of only 11.3%, compared with 54.4% for the monovalent injected vaccine in Asian seabass ^78, 51^and a bivalent *F. oreochromis* and *F. orientalis* vaccine similarly declined from 50% to 38.89% ^49^. Furthermore, polyvalent vaccines incorporating multiple CCB species are currently absent from the literature, representing a gap that future research on broadly protective vaccines could address.

Several aspects of trial design may have further influenced the reliability of individual efficacy estimates. The virulence of the challenge strain, for example, can directly affect measured protection. Several trials used the *F. psychrophilum* NCIMB 1947ᵀ strain for challenge ^79^, which is weakly virulent and has limited biofilm-forming capacity, potentially underestimating vaccine efficacy ^80,81^. The results of these trials should therefore be interpreted with caution, reinforcing the need for standardized challenge models that use characterised strains of confirmed virulence. In addition, more than half of the comparisons (56.7%) were rated as having “some concerns” for the randomisation process, which may have systematically inflated efficacy estimates. Publication bias may also have contributed: the funnel plot was asymmetric and Egger’s test was significant (*P* <0.001), suggesting that studies reporting low or null efficacy are underrepresented and that the overall efficacy may have been overestimated. Sensitivity analyses showed that the pooled RR remained stable (0.54–0.57) after excluding trials with 100% control mortality, those with small sample sizes (n <50 per group). This indicates that the main finding is not dependent on any particular subset of trials or extreme experimental conditions. The consistently high heterogeneity across all sensitivity models (*I*² >89%) further confirms that variation in vaccine efficacy reflects biological and methodological options, rather than being driven by a few influential comparisons. Several limitations of this study should be acknowledged. First, the efficacy of conventional vaccines (inactivated and live attenuated) and modern nanoparticle-based delivery platforms (e.g., chitosan, cationic lipid nanoparticles) was not compared separately; rather, they were grouped into broad vaccine categories, which may have inflated the observed heterogeneity. Second, some unmeasured variables, including adjuvant type, challenge dose, booster interval, and vaccine concentration, could not be incorporated into subgroup analyses and may have further contributed to the heterogeneity. Third, the number of comparisons was limited for certain subgroups (*F. johnsoniae*, *k* = 4; *F. davisii* =10), and the resulting wide confidence intervals restrict the reliability of inferences. Fourth, the search was limited to English-language publications, which may have introduced language-based bias.

Taken together, these findings point to several priorities for future research. First, *F. oreochromis* and *F. columnare* should be treated as priority targets for vaccine development: both showed relatively low protective efficacy (RPS = 36.3% and 30.6%, respectively), with *F. columnare* being the lowest. Moreover, *F. columnare* also has the highest intrinsic virulence (median control mortality 90%) and exhibits asymmetric cross-protective capacity. Second, although *F. columnare* is known to infect salmonids and cause columnaris disease ^82,83^, all eligible vaccine challenge trials published between years 2000 and 2026 were conducted in grass carp. Early attempts to evaluate *F. columnare* vaccines in salmonids were reported before year 2000, but most showed limited efficacy ^84,85^. Notably, the past two decades have seen no experimental challenge trials evaluating *F. columnare* vaccines in salmonids, leaving a gap in the evidence base. A recent review similarly noted that vaccine candidates against diseases caused by *F. columnare* and *F. davisii* remain almost entirely unevaluated in salmonid species, which underscores the urgency of vaccine efficacy evaluation against *F. columnare* in salmonid hosts ^2^. Third, *F. branchiophilum*, the primary agent of bacterial gill disease ^86,87^, is not represented by any vaccine study meeting our inclusion criteria after year 2000 ^88,89^, highlighting an imbalance in pathogen coverage within *Flavobacterium* vaccine research. Finally, improvements in study design are equally critical. The adoption of standardized challenge models and more rigorous reporting of randomization and blinding would enhance cross-study comparability and provide a more reliable evidence base for future quantitative syntheses.

## 5 Conclusion

This systematic review and meta-analysis demonstrate that vaccination provides substantial protection against *Flavobacterium* infections in farmed fish, with an overall pooled RPS of 42.8%. However, extensive heterogeneity was observed in this overall estimate, driven by complex interactions among pathogen species, fish host and vaccine type. Live attenuated vaccines provided the highest protection, followed by inactivated vaccines, while subunit vaccines showed substantially lower efficacy. Cross-protection was highly asymmetric, with *F. covae*-derived vaccines showing broader protection activity, a property that was further influenced by host association. Polyvalent vaccines were effective against *F. psychrophilum* but consistently failed to improve protection against CCB, and polyvalent formulations incorporating the CCB group are currently lacking. Evidence of publication bias and inconsistent reporting of blinding suggests that the overall efficacy may be overestimated. Future vaccine development should prioritize *F. oreochromis* and *F. columnare*, address this evidence gap for *F. columnare* in salmonids, and adopt standardized challenge models with more rigorous reporting of randomization and blinding to improve cross-study comparability.

## Acknowledgements

This study was supported by the APRC-CityU New Research Initiatives/Infrastructure Support from Central (9610574, 7006064), and Early career scheme (project number 21103324) from the Hong Kong Research Grant Council.

## Appendices

**Appendix I.** Supplementary Table S1. Summary of all included studies.

**Appendix II**. Supplementary Table S2. Detailed results for *F. psychrophilum* subgroup analyses.

**Appendix III**. Supplementary Table S3. Cross–protection and polyvalent vaccine efficacy data.

**Appendix IV**. Supplementary Figure S1. Funnel plot for publication bias.

**Appendix V**. Supplementary Figure S2. Risk of bias assessment details.

**Appendix VI**. Supplementary Figure S3. Leave–one–study–out sensitivity analysis.

**Appendix VII**. Supplementary Figure S4. Cross–protection among Flavobacterium species.

**Appendix VIII**. Supplementary File S1. Search strings used in PubMed, Web of Science, and Scopus.

