## Supplementary figures and images for "Efficacy of vaccines against *Flavobacterium* infections in fish: A systematic review and meta-analysis"

### Figure S1

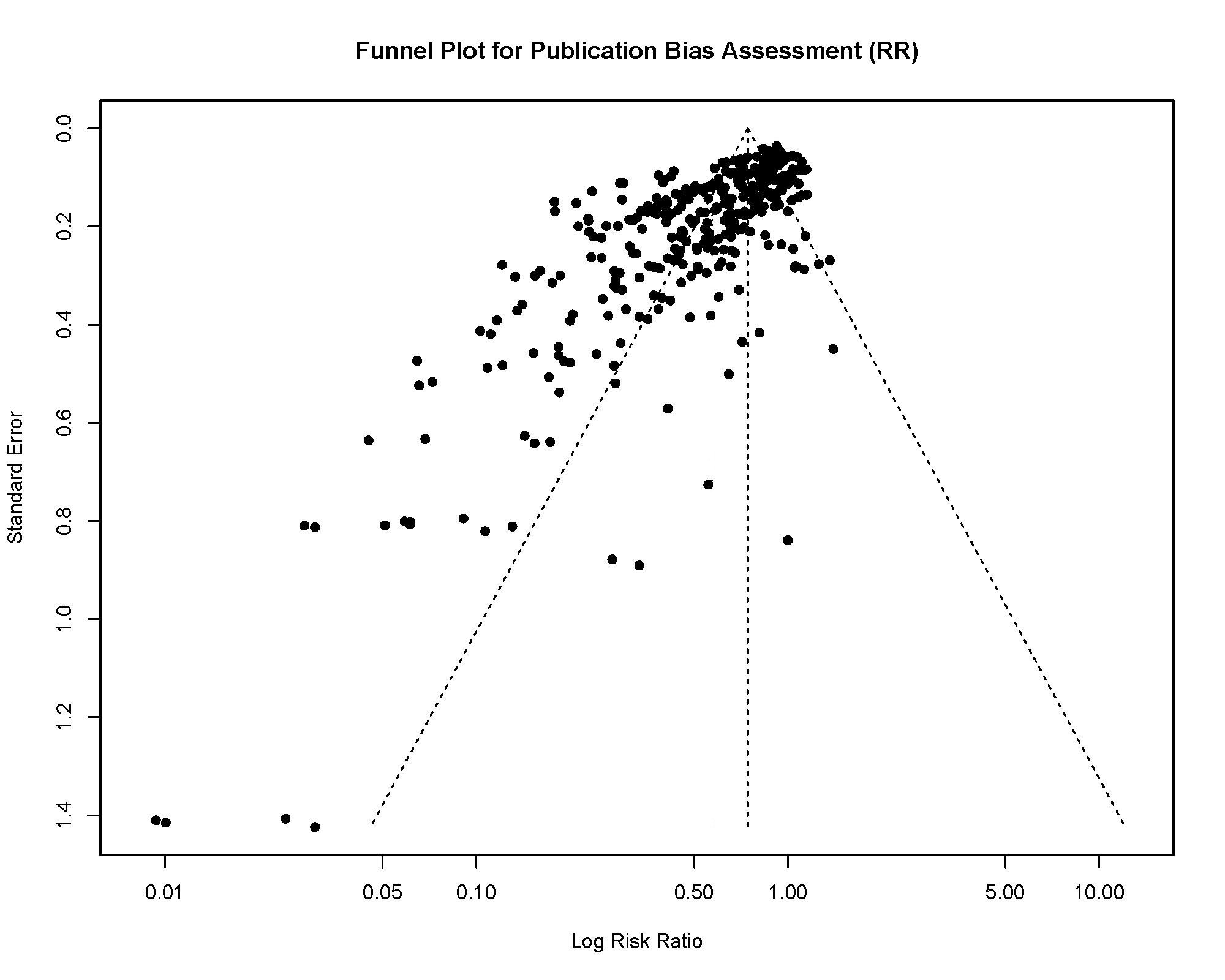

### Figure S2

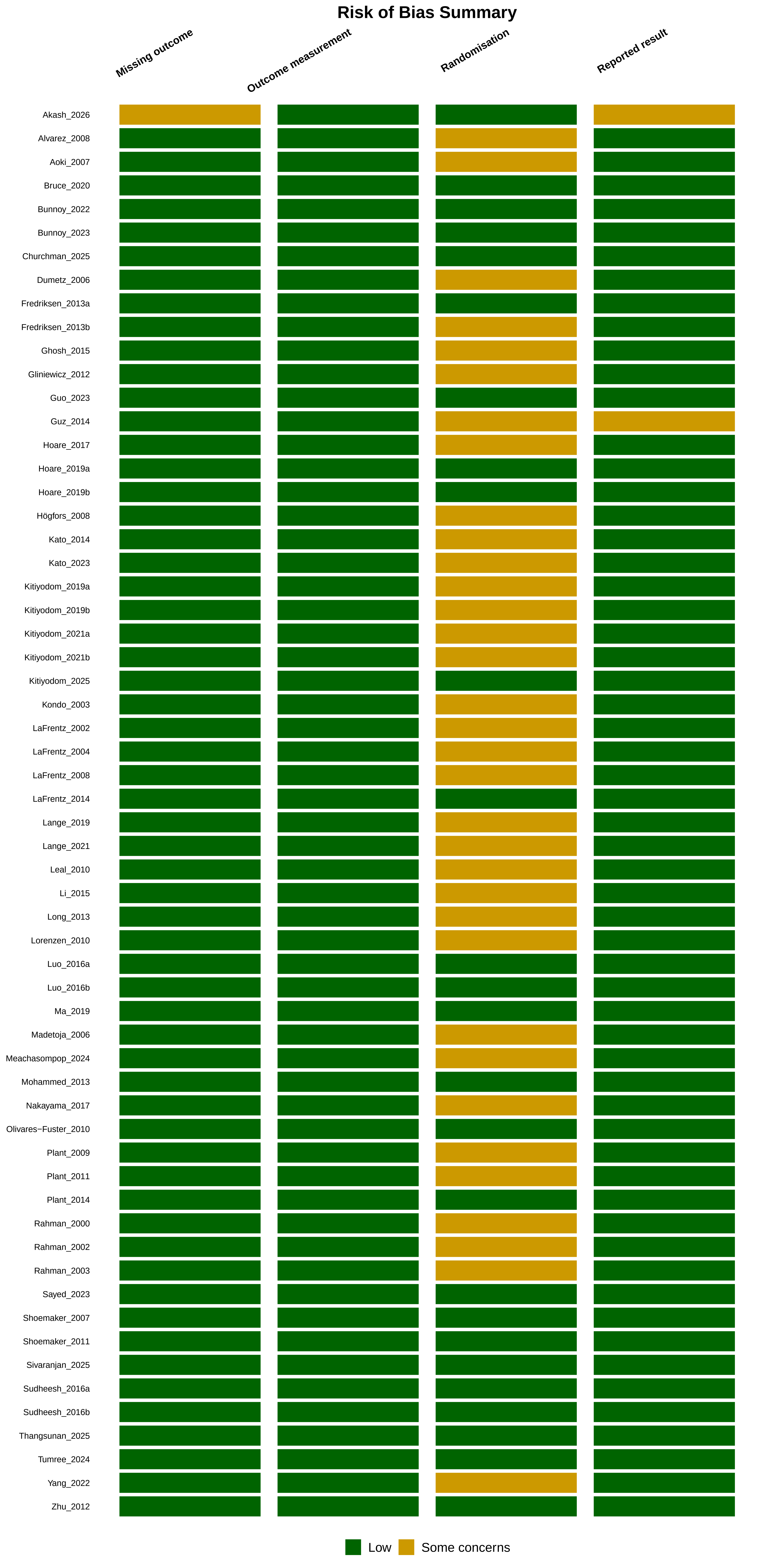

### Figure S4

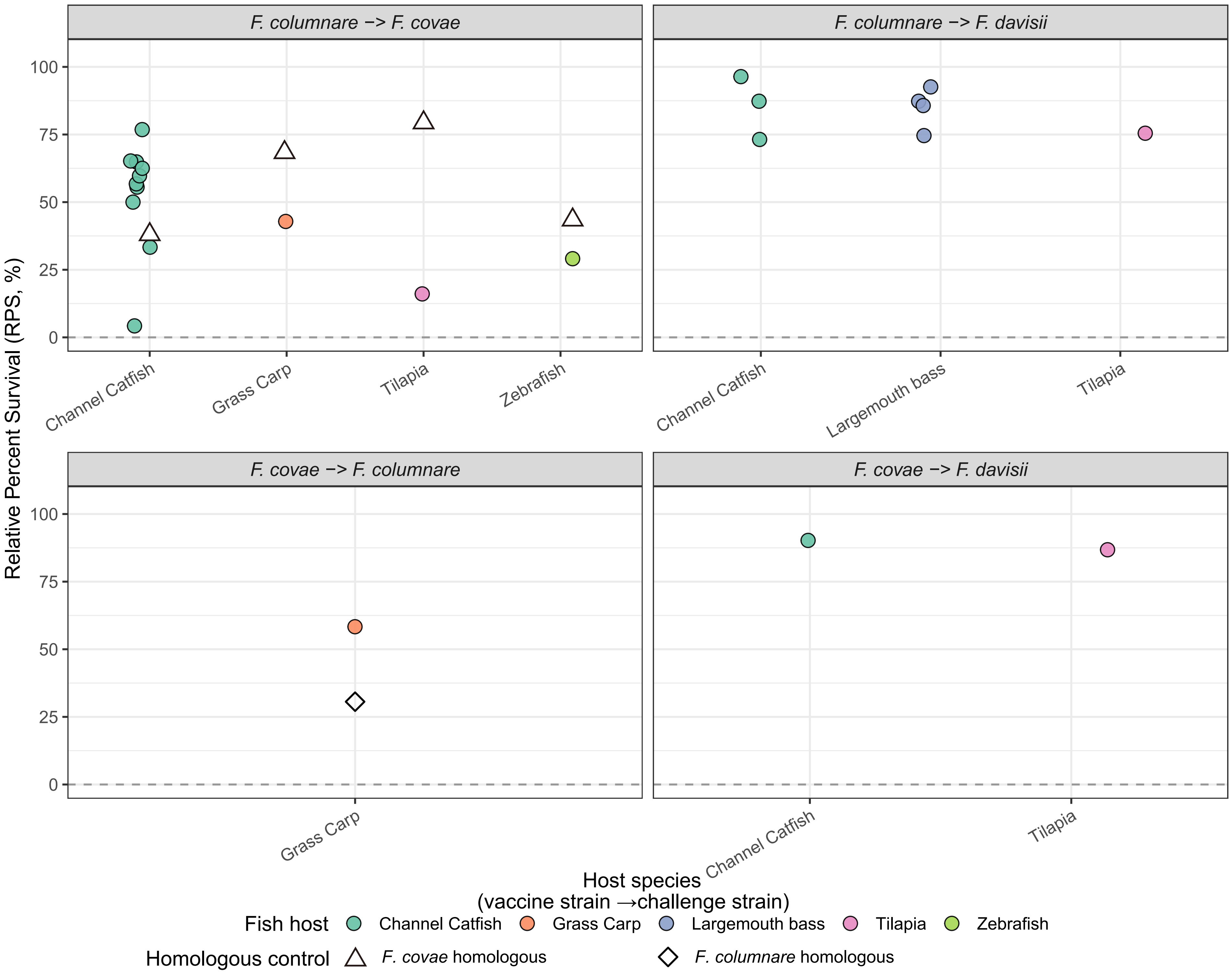
