## Supplementary material for "Efficacy of vaccines against *Flavobacterium* infections in fish: A systematic review and meta-analysis": Figure S3

### Leave−one−study−out Sensitivity Analysis (RR)

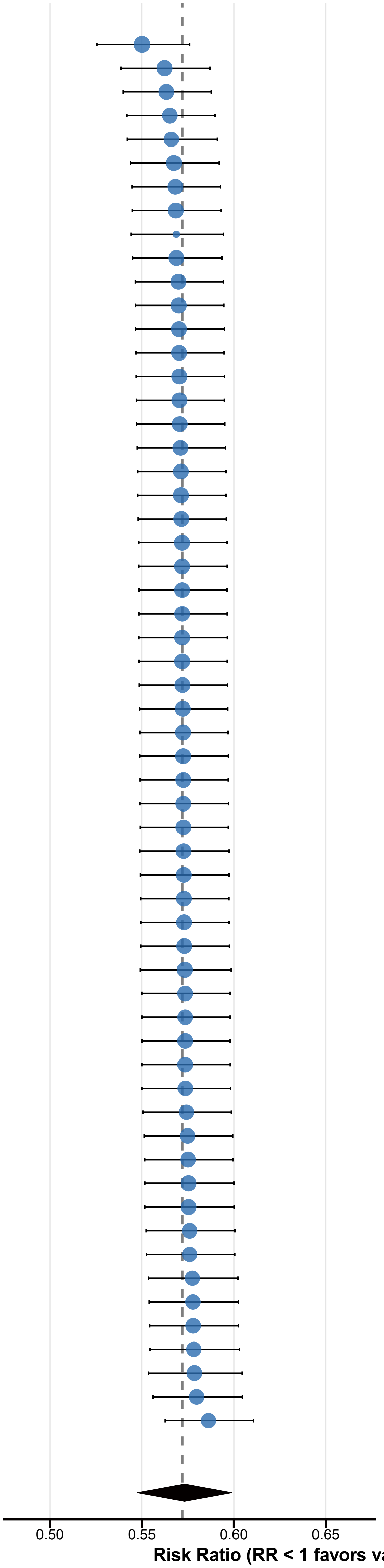


| Study | Pooled RR [95% CI] | I² |
|---|---|---|
| Guo_2023 | 0.550 [0.525, 0.576] | 90.1% |
| Plant_2014 | 0.562 [0.539, 0.587] | 89.8% |
| Leal_2010 | 0.563 [0.540, 0.588] | 89.6% |
| Plant_2009 | 0.565 [0.542, 0.590] | 89.5% |
| Ghosh_2015 | 0.566 [0.542, 0.591] | 89.4% |
| Plant_2011 | 0.567 [0.544, 0.592] | 89.7% |
| Guz_2014 | 0.568 [0.545, 0.593] | 89.6% |
| LaFrentz_2008 | 0.568 [0.545, 0.593] | 89.7% |
| Kitiyodom_2025 | 0.569 [0.544, 0.594] | 87.2% |
| Lange_2021 | 0.569 [0.545, 0.594] | 89.6% |
| Lorenzen_2010 | 0.570 [0.546, 0.594] | 89.5% |
| Tumree_2024 | 0.570 [0.546, 0.595] | 89.6% |
| Bunnoy_2022 | 0.570 [0.546, 0.595] | 89.5% |
| Olivares−Fuster_2010 | 0.570 [0.547, 0.595] | 89.5% |
| Kato_2023 | 0.570 [0.547, 0.595] | 89.6% |
| Gliniewicz_2012 | 0.570 [0.547, 0.595] | 89.6% |
| LaFrentz_2014 | 0.571 [0.547, 0.595] | 89.6% |
| Yang_2022 | 0.571 [0.547, 0.596] | 89.6% |
| Kitiyodom_2019a | 0.571 [0.548, 0.596] | 89.5% |
| Kato_2014 | 0.571 [0.548, 0.596] | 89.6% |
| Thangsunan_2025 | 0.571 [0.548, 0.596] | 89.6% |
| Meachasompop_2024 | 0.572 [0.548, 0.596] | 89.6% |
| Churchman_2025 | 0.572 [0.548, 0.596] | 89.6% |
| Luo_2016b | 0.572 [0.548, 0.596] | 89.5% |
| Li_2015 | 0.572 [0.548, 0.596] | 89.5% |
| Sayed_2023 | 0.572 [0.548, 0.597] | 89.6% |
| Rahman_2000 | 0.572 [0.548, 0.596] | 89.6% |
| Bunnoy_2023 | 0.572 [0.549, 0.597] | 89.6% |
| Dumetz_2006 | 0.572 [0.549, 0.597] | 89.6% |
| Nakayama_2017 | 0.572 [0.549, 0.597] | 89.6% |
| Lange_2019 | 0.572 [0.549, 0.597] | 89.6% |
| Fredriksen_2013b | 0.573 [0.549, 0.597] | 89.5% |
| LaFrentz_2002 | 0.573 [0.549, 0.597] | 89.5% |
| Bruce_2020 | 0.573 [0.549, 0.597] | 89.5% |
| LaFrentz_2004 | 0.573 [0.549, 0.597] | 89.5% |
| Luo_2016a | 0.573 [0.549, 0.597] | 89.6% |
| Hoare_2017 | 0.573 [0.549, 0.597] | 89.5% |
| Álvarez_2008 | 0.573 [0.549, 0.597] | 89.5% |
| Sudheesh_2016b | 0.573 [0.549, 0.598] | 89.5% |
| Mohammed_2013 | 0.573 [0.549, 0.599] | 89.7% |
| Kitiyodom_2021a | 0.574 [0.550, 0.598] | 89.5% |
| Zhu_2012 | 0.574 [0.550, 0.598] | 89.4% |
| Högfors_2008 | 0.574 [0.550, 0.598] | 89.6% |
| Hoare_2019b | 0.574 [0.550, 0.598] | 89.6% |
| Aoki_2007 | 0.574 [0.550, 0.598] | 89.5% |
| Kitiyodom_2021b | 0.574 [0.551, 0.599] | 89.5% |
| Sivaranjan_2025 | 0.575 [0.551, 0.599] | 89.5% |
| Hoare_2019a | 0.575 [0.552, 0.600] | 89.5% |
| Rahman_2002 | 0.575 [0.552, 0.600] | 89.7% |
| Rahman_2003 | 0.575 [0.552, 0.600] | 89.7% |
| Long_2013 | 0.576 [0.552, 0.601] | 89.5% |
| Fredriksen_2013a | 0.576 [0.553, 0.600] | 89.3% |
| Shoemaker_2007 | 0.577 [0.554, 0.602] | 89.4% |
| Kondo_2003 | 0.578 [0.554, 0.603] | 89.5% |
| Shoemaker_2011 | 0.578 [0.554, 0.603] | 89.5% |
| Kitiyodom_2019b | 0.578 [0.554, 0.603] | 89.4% |
| Sudheesh_2016a | 0.579 [0.554, 0.605] | 89.6% |
| Ma_2019 | 0.580 [0.556, 0.605] | 89.4% |
| Madetoja_2006 | 0.586 [0.563, 0.611] | 89.2% |
| **Overall** | 0.572 [0.549, 0.597] | |
