## Supplementary material for "Efficacy of vaccines against *Flavobacterium* infections in fish: A systematic review and meta-analysis": File S1

**Supplementary materials**

**S1 Search String:**

**PubMed**
((("fishes"[MeSH Terms] OR "Salmonidae"[MeSH Terms] OR "Oncorhynchus mykiss"[MeSH Terms] OR "Tilapia"[MeSH Terms] OR "Carps"[MeSH Terms] OR "Ictaluridae"[MeSH Terms] OR "Eels"[MeSH Terms] OR "Perches"[MeSH Terms] OR "bass"[MeSH Terms] OR "fish*"[Title/Abstract] OR "finfish*"[Title/Abstract] OR "teleost*"[Title/Abstract] OR "ichthyo*"[Title/Abstract] OR "pisciculture*"[Title/Abstract] OR "salmon*"[Title/Abstract] OR "trout*"[Title/Abstract] OR "oncorhynchus*"[Title/Abstract] OR "tilapia*"[Title/Abstract] OR "oreochromis*"[Title/Abstract] OR "carp*"[Title/Abstract] OR "cyprinid*"[Title/Abstract] OR "catfish*"[Title/Abstract] OR "ictalurus*"[Title/Abstract] OR "seabass"[Title/Abstract] OR "sea bass"[Title/Abstract] OR "dicentrarchus"[Title/Abstract] OR "lateolabrax"[Title/Abstract] OR "large yellow croaker"[Title/Abstract] OR "larimichthys*"[Title/Abstract] OR "eel"[Title/Abstract] OR "anguilla*"[Title/Abstract]) AND ("Vaccines"[MeSH Terms] OR "Vaccination"[MeSH Terms] OR "Immunization"[MeSH Terms] OR "vaccin*"[Title/Abstract] OR "immuni*"[Title/Abstract] OR "immunoprophyla*"[Title/Abstract] OR "bacterin*"[Title/Abstract] OR "killed vaccine*"[Title/Abstract] OR "inactivated"[Title/Abstract] OR "live attenuated"[Title/Abstract] OR "attenuated"[Title/Abstract] OR "subunit vaccine*"[Title/Abstract] OR "recombinant vaccine*"[Title/Abstract] OR "dna vaccine*"[Title/Abstract] OR "genetic vaccine*"[Title/Abstract] OR "plasmid*"[Title/Abstract]) AND ("Survival Rate"[MeSH Terms] OR "Mortality"[MeSH Terms] OR "challenge*"[Title/Abstract] OR "experimental infection*"[Title/Abstract] OR "artificial infection*"[Title/Abstract] OR "relative percent survival"[Title/Abstract] OR "RPS"[Title/Abstract] OR "survival*"[Title/Abstract] OR "mortalit*"[Title/Abstract] OR "protect*"[Title/Abstract] OR "efficac*"[Title/Abstract] OR "effectiveness"[Title/Abstract]) AND ("Flavobacterium"[MeSH Terms] OR "flavobacterium*"[Title/Abstract] OR "flavobacteri*"[Title/Abstract] OR "columnari*"[Title/Abstract] OR "Coldwater disease"[Title/Abstract] OR "CWD"[Title/Abstract] OR "Rainbow trout fry syndrome"[Title/Abstract] OR "RTFS"[Title/Abstract] OR "Bacterial gill disease"[Title/Abstract] OR "BGD"[Title/Abstract] OR "Cytophaga psychrophila"[Title/Abstract])) NOT "REVIEW"[Publication Type]) AND 2000/01/01:2026/03/31[Date - Publication] AND "English"[Language]

**Web of Science**

(TS=(fish* OR finfish* OR teleost* OR ichthyo* OR pisciculture* OR salmon* OR trout* OR oncorhynchus* OR tilapia* OR oreochromis* OR carp* OR cyprinid* OR catfish* OR ictalurus* OR seabass OR "sea bass" OR dicentrarchus OR lateolabrax OR "large yellow croaker" OR larimichthys* OR eel* OR anguilla*) AND TS=(vaccin* OR immuni* OR immunoprophyla* OR bacterin* OR "killed vaccine*" OR inactivated OR "live attenuated" OR attenuated OR "subunit vaccine*" OR "recombinant vaccine*" OR "DNA vaccine*" OR "genetic vaccine*" OR plasmid*) AND TS=(challenge* OR "experimental infection*" OR "artificial infection*" OR "relative percent survival" OR RPS OR survival* OR mortalit* OR protect* OR efficac* OR effectiveness) AND TS=(Flavobacterium* OR flavobacteri* OR columnari* OR "Coldwater disease" OR CWD OR "Rainbow trout fry syndrome" OR RTFS OR "Bacterial gill disease" OR BGD OR "Cytophaga psychrophila")) AND LA=(English OR Unspecified) NOT PY=(1968 OR 1969 OR 1976 OR 1977 OR 1981 OR 1982 OR 1983 OR 1986 OR 1987 OR 1992 OR 1994 OR 1995 OR 1997 OR 1999) NOT DT=(Book OR Meeting OR Abstract OR Awarded Grant OR Review Article OR Editorial Material OR Correction OR Retracted Publication)

**Scopus**

TITLE-ABS-KEY( (fish* OR finfish* OR teleost* OR ichthyo* OR pisciculture* OR salmon* OR trout* OR oncorhynchus* OR tilapia* OR oreochromis* OR carp* OR cyprinid* OR catfish* OR ictalurus* OR seabass OR "sea bass" OR dicentrarchus OR lateolabrax OR "large yellow croaker" OR larimichthys* OR eel* OR anguilla*) AND (vaccin* OR immuni* OR immunoprophyla* OR bacterin* OR "killed vaccine*" OR inactivated OR "live attenuated" OR attenuated OR "subunit vaccine*" OR "recombinant vaccine*" OR "DNA vaccine*" OR "genetic vaccine*" OR plasmid*) AND (challenge* OR "experimental infection*" OR "artificial infection*" OR "relative percent survival" OR RPS OR survival* OR mortalit* OR protect* OR efficac* OR effectiveness) AND (Flavobacterium* OR flavobacteri* OR "Coldwater disease" OR CWD OR "Rainbow trout fry syndrome" OR RTFS OR columnari* OR "Bacterial gill disease" OR BGD OR "Cytophaga psychrophila"))AND PUBYEAR > 1999 AND LANGUAGE(english) AND NOT (DOCTYPE(re) OR DOCTYPE(ed) OR DOCTYPE(ch))
